# Metagenomic characterization of creek sediment microbial communities from a major agricultural region in Salinas, California

**DOI:** 10.1101/737759

**Authors:** Brittany J. Suttner, Eric R. Johnston, Luis H. Orellana, Luis M. Rodriguez-R, Janet K. Hatt, Diana Carychao, Michelle Q. Carter, Michael B. Cooley, Konstantinos T. Konstantinidis

## Abstract

Little is known about the public health risks associated with natural creek sediments that are affected by runoff and fecal pollution from agricultural and livestock practices. For instance, the persistence of foodborne pathogens originating from agricultural activities such as Shiga Toxin-producing *E. coli* (STEC) in such sediments remains poorly quantified. Towards closing these knowledge gaps, the water-sediment interface of two creeks in the Salinas River Valley was sampled over a nine-month period using metagenomics and traditional culture-based tests for STEC. Our results revealed that these sediment communities are extremely diverse and comparable to the functional and taxonomic diversity observed in soils. With our sequencing effort (~4 Gbp per library), we were unable to detect any pathogenic *Escherichia coli* in the metagenomes of 11 samples that had tested positive using culture-based methods, apparently due to relatively low pathogen abundance. Further, no significant differences were detected in the abundance of human- or cow-specific gut microbiome sequences compared to upstream, more pristine (control) sites, indicating natural dilution of anthropogenic inputs. Notably, a high baseline level of metagenomic reads encoding antibiotic resistance genes (ARGs) was found in all samples and was significantly higher compared to ARG reads in metagenomes from other environments, suggesting that these communities may be natural reservoirs of ARGs. Overall, our metagenomic results revealed that creek sediments are not a major sink for anthropogenic runoff and the public health risk associated with these sediment microbial communities may be low.

**IMPORTANCE:** Current agricultural and livestock practices contribute to fecal contamination in the environment and the spread of food and water-borne disease and antibiotic resistance genes (ARGs). Traditionally, the level of pollution and risk to public health is assessed by culture-based tests for the intestinal bacterium, *E. coli*. However, the accuracy of these traditional methods (e.g., low quantification, and false positive signal when PCR-based) and their suitability for sediments remains unclear. We collected sediments for a time series metagenomics study from one of the most highly productive agricultural regions in the U.S. in order to assess how agricultural runoff affects the native microbial communities and if the presence of STEC in sediment samples can be detected directly by sequencing. Our study provided important information on the potential for using metagenomics as a tool for assessment of public health risk in natural environments.

## INTRODUCTION

Nearly half of the major produce-associated outbreaks in the U.S. between 1995-2006 have been traced to spinach or lettuce grown in the Salinas Valley of California (1). Contamination of produce can be caused by exposure to contaminated irrigation or flood water, deposition of feces by wildlife or livestock, or during field application of manure as fertilizer (2, 3). From a public health perspective, more information is needed on the risk of exposure to animal fecal contamination as recent studies suggest that exposure to water impacted by cow feces may present public health risks that are similar or equal to human fecal contamination. For example, cattle are a reservoir of the major foodborne pathogen, Shiga Toxin-producing *E. coli* (STEC) (4, 5). Environmental contamination by animal feces from farms is an emerging public health issue not only as a source of pathogens but also as a source of antibiotic resistance genes (ARGs) (6). Antibiotics are regularly administered to livestock at prophylactic concentrations to prevent infection, and food animal production is responsible for a significant proportion of total antibiotic use (7). Such practices are known to contribute to the prevalence of ARGs in the environment (8–10), which can spread rapidly to other microbes via horizontal gene transfer, including to human pathogens of clinical importance (11, 12). Surprisingly, there is very little regulation of antibiotic use in the livestock industry, even though these operations can be major contributors to fecal pollution and the spread of ARGs in the environment (13, 14).

Our previous culture- and PCR-based surveys of the Salinas watershed, and particularly Gabilan and Towne Creeks (heretofore called GABOSR and TOWOSR, respectively), indicated persistent presence of STEC in water and sediments (15, 16) and a potentially significant public health risk. Continued prevalence of STEC in both GABOSR and TOWOSR sites is hypothesized to be linked to the presence of cattle upstream. For instance, in several cases, STEC strains isolated from cattle fecal samples were identical to those found in water and sediment based on Multi-Locus Variable number tandem repeat Analysis (MLVA) typing. Indeed, the prevalence of STEC was strongly correlated with runoff due to rainfall (1, 16). However, hydrologic modeling and surveys indicated that pathogen levels in streams were not only due to overland flow, but also to contributions from sediment (17, 18). These observations were further supported by several examples of identical MLVA types isolated from both water and sediment at the same location or downstream during periods of drought (1, 15). Further, the levels of pathogen in the water column and sediment are difficult to measure and are generally underestimated due to the predominance of biofilms and viable but not culturable (VBNC) bacteria (19). Therefore, metagenomic characterization of the creek sediments should provide independent quantitative insights into the effect of agricultural practices on the surrounding environment.

River and creek sediments are among the most diverse communities sequenced to date and are largely under-sampled (20, 21). Moreover, the sediments studied to date are exclusively from highly and/or historically polluted environments with varying industrial or sewage inputs and thus, each sediment is characterized by its unique properties in terms of flow dynamics, chemical environment, climatic conditions and anthropogenic inputs (21–27). Accordingly, previous studies on the effect of anthropogenic inputs on sediments in lotic (free-flowing) aquatic systems have yielded mixed results on how surrounding land use practices impact sediment communities or were not directly relevant. Furthermore, in order to properly quantify the effect of anthropogenic antibiotic inputs, appropriate controls (e.g., pristine sampling sites) are needed to determine baseline levels of ARGs and other genes (13, 28).

In this study, we examined the effect of agricultural runoff on microbial communities from creek sediments in the Salinas watershed and whether community structure correlated with precipitation or culture-based detection of STEC. We sampled nearby, upstream sites with reduced human and cattle presence as a baseline to compare the abundance of anthropogenic signals (i.e. human and cow gut microbiome and ARGs) observed in the downstream sites. Furthermore, we compared these sites to other publicly-available sediment, soil, and river water metagenomes from both highly pristine and polluted environments in order to validate our results and assess anthropogenic pollution levels relative to other similar habitats.

## RESULTS

### Description of sampling sites

Six sites from three creeks in the Salinas River valley in California were included in this study. Two of the sites (collectively referred to as the “downstream” samples/sites) are impacted by cattle ranching but vary in the level of agricultural activities in the directly surrounding area. The creeks are isolated at the sampling locations but converge further downstream before emptying into the Salinas River. Gabilan (GABOSR) is directly downstream of organic strawberry produce fields that use both green and poultry manure fertilizer and has cattle ranching upstream of the strawberry farm. The second site, Towne Creek (TOWOSR), is roughly 2 Km north of GABOSR but does not have any abutting agricultural fields directly upstream and only receives input from cattle ranches. Ten samples from each of the two downstream sites, GABOSR and TOWOSR, collected over a 9-month period from September 2013 through June 2014 were selected for metagenome sequencing based on precipitation levels and detection of pathogenic *E. coli* via enrichment culture (Table 1). An additional seven samples from four upstream sites (collectively referred to as the “upstream” samples/sites), were included to serve as upstream controls for metagenomic comparison (Table 1 and Figure 1). The samples from these locations included: three samples collected ~10 km upstream from Gabilan (“GABOSR Control”) on March 2016 (GC1-3); two samples collected ~3 km upstream from Towne Creek (“TOWOSR Control”) on April 2017 (TC1 and TC2); and finally, one sample from each of two sites on the west side of the Salinas River (“West Salinas”), ~60 km and 110 km southeast from the downstream sites collected in May 2017 (WS1 and WS2, respectively). The latter two samples are not upstream of GABOSR or TOWOSR but were included because they are more pristine sites with no known history of cattle impact, as opposed to the GC and TC samples, which may have had minimal inputs from previous cattle grazing.

**Table 1:** Culture-based detection of STEC and precipitation (Precip) data reported in inches

| Sample ID | Date Collected | STEC<sup>a</sup> | Copies *stx2*/ug DNA<sup>b</sup> | Precip | 5-day Precip |
| --- | --- | --- | --- | --- | --- |
| **Gabilan at Old Stage (GABOSR)** |  |  |  |  |  |
| G130904 | 9/4/13 | - | 8.1 | 0 | 0 |
| G140116 | 1/16/14 | + | 8 | 0 | 0.01 |
| G140128 | 1/28/14 | + | 0 | 0 | 0 |
| G140210 | 2/10/14 | + | 4.4 | 0.01 | 1.1 |
| G140224 | 2/24/14 | + | 1.8 | 0 | 0 |
| G140301 | 3/1/14 | + | 1.5 | 0.33 | 2.01 |
| G140319 | 3/19/14 | - | 0 | 0 | 0.01 |
| G140402 | 4/2/14 | + | 1.4 | 0.03 | 1.04 |
| G140415 | 4/15/14 | - | 0 | 0 | 0 |
| G140611 | 6/11/14 | - | 2.4 | 0 | 0 |
| **Towne Creek at Old Stage (TOWOSR)** |  |  |  |  |  |
| T130904 | 9/4/13 | - | 14.2 | 0 | 0 |
| T130918 | 9/18/13 | + | 15.3 | 0 | 0 |
| T131023 | 10/23/13 | - | 0 | 0 | 0 |
| T131230 | 12/30/13 | + | 3.9 | 0 | 0 |
| T140116 | 1/16/14 | - | 0 | 0 | 0.01 |
| T140128 | 1/28/14 | + | 0 | 0 | 0 |
| T140210 | 2/10/14 | - | 1.7 | 0.01 | 1.1 |
| T140224 | 2/24/14 | - | 1.5 | 0 | 0 |
| T140319 | 3/19/14 | - | 0 | 0 | 0.01 |
| T140611 | 6/11/14 | - | 0 | 0 | 0 |
| **Upstream GABOSR Control (GC)** |  |  |  |  |  |
| GC1 | 3/9/16 | - | 0 | 0 | 2.84 |
| GC2 | 3/9/16 | + | 0 | 0 | 2.84 |
| GC3 | 3/9/16 | - | 0 | 0 | 2.84 |
| **Upstream TOWOSR Control (TC)** |  |  |  |  |  |
| TC1 | 4/19/17 | + | 0 | 0 | 0.45 |
| TC2 | 4/19/17 | - | 0 | 0 | 0.45 |
| **West Salinas (WS)** |  |  |  |  |  |
| WS1 | 5/4/17 | - | 0 | 0 | 0 |
| WS2 | 5/4/17 | - | 0 | 0 | 0 |

<sup>a</sup> Samples in which STEC was detected by PCR of enrichment cultures are listed as either positive (+) or negative (-).
<sup>b</sup> Copy number of the shiga toxin gene (*stx2*) was determined via ddPCR.

**Figure 1:**
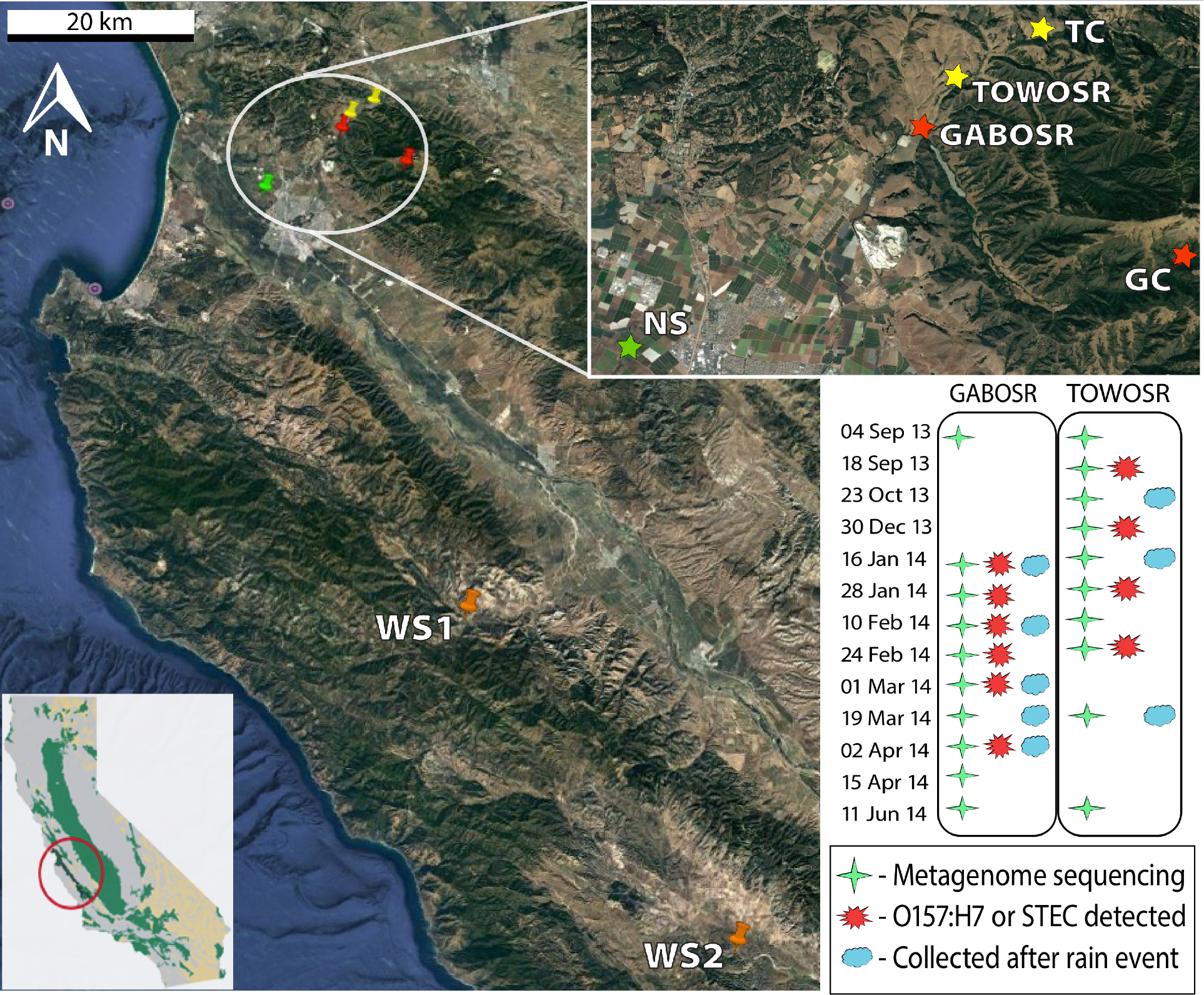
Location of sampling sites in the Salinas Valley, California and sampling scheme for time-series metagenomics. Sampling site for Gabilan (GABOSR in red) and Towne Creek (TOWOSR in yellow). The upstream controls for Gabilan (GC) and Towne Creek (TC) are also indicated by the same colors. Orange pins mark the West Salinas sites (WS1 and WS2) included as less agriculturally-impacted controls. The North Salinas weather station (NS; green star) is approximately 11km SE of GABOSR and was the closest weather monitoring station to all samples shown in the subset map. GPS coordinates for all sampling locations are provided in supplementary Table S1. Inset: location of the Salinas Valley in the state of California.

### Description of metagenomes and sequence coverage of microbial community

A total of 27 metagenomic samples, ranging in size from 8.7 to 20.1 million reads (2.5 to 5 Gbp) after trimming, were recovered from the six locations (Table S2). For all samples, less than 28% of the total community (average 18.6%) was covered by our sequencing efforts as determined by Nonpareil analysis (Figure S1). Consequently, the assembly of the metagenomes was limiting (Table S2), consistent with our previous analysis of soil and sediment communities (29) and those of a few other metagenomic studies of river sediments. Thus, an un-assembled short read-based strategy was used for all subsequent analyses (paired-end, non-overlapping reads with an average length of 132-145 bp per dataset), unless noted otherwise. A total of 7.2×10^8^ protein sequences were predicted from the short reads, with an average of 2.7×10^7^ sequences per sample. The number of protein sequences that could be annotated to the Swiss-Prot database in each sample ranged between 10 and 16% (average 14.5%) of the total sequences.

### OTU characterization and alpha diversity assessment

A total of 466,421 reads encoding fragments of the 16S or 18S rRNA gene were detected in all 27 metagenomes with an average of 17,275 reads per sample. All datasets were dominated by bacteria, with only 0.6% and 3.0% of the total rRNA reads, on average, having archaeal or eukaryotic origin, respectively. Closed-reference OTU picking at 97% nucleotide identity threshold resulted in a total of 25,764 OTUs from 349,886 reads for all 27 samples and an average of 4,465 OTUs per sample. Since the coverage was similar for all datasets, the number of OTUs shared between all samples were compared without any further normalization. Only 138 OTUs (0.5%) were shared among all 27 samples, while 9,500 (36.9%) of the OTUs were present in only one sample. The OTU rarefaction plot showed that diversity was not saturated (Figure S2A), which agreed with the low number of shared OTUs and the Nonpareil estimates on the shotgun data reported above (Figure S1).

Alpha diversity observed in the California samples was compared to three publicly-available river sediment metagenomes from Montana that had similar land use inputs (i.e. agricultural or small towns) and were the most appropriate data for comparison among lotic sediment metagenomes currently available (20). Species richness and diversity in Montana samples were significantly less than California samples (P= 2.3×10^−4^ and 0.006, respectively; Figure S2). Within California sites, diversity and evenness were similar; however, average species richness in GABOSR was significantly lower than TOWOSR and the upstream samples (P= 0.034 and 4.1×10^−4^, respectively).

### Taxonomic composition and functional diversity of water-sediment microbial communities

OTUs were analyzed further to characterize the taxonomic profile of the communities sampled. *Proteobacteria* and *Bacteroidetes* were the most abundant phyla across most samples. However, some of the upstream samples had a higher abundance of *Actinobacteria* (Figure S3A). Class level taxonomic distributions were consistent over time for GABOSR samples and revealed the high abundance of *Betaproteobacteria* (>19-24% of total sequences). TOWOSR samples varied more over time; five samples (T130918,T131230, T140128, T140210, T140611) had a higher abundance of *Deltaproteobacteria* and *Bacteroidia*, and one sample (T140116) had a higher abundance of *Cyanobacteria*. The upstream samples also showed a similar community composition and had higher relative abundance of *Alphaproteobacteria* (11-17%) compared to the downstream samples (Figure S3B). These results were consistent with the TrEMBL taxonomic classification of protein-coding metagenomic reads, which were dominated by *Bacteria* (~95.2% per sample; Figure S4).

### Microbial community structure and dynamics in Salinas River valley creeks

Location was the strongest factor affecting clustering patterns observed in PCA ordinations of all distance matrices analyzed (Figure S5). ADONIS analysis in the R package vegan (using location as a categorical variable) yielded *P*<0.001 and R^2^= 0.44, 0.67, 0.41, and 0.56 for MASH, functional gene, OTUs Bray-Curtis (16S-BC) and OTUs weighted UniFrac (16S-WUF), respectively. This result was confirmed by correlation analysis of the NMDS ordinations to all metadata variables using the envfit function in vegan. After Bonferroni correction for multiple comparisons, location had the strongest correlation to all ordinations (MASH: P=0.001, R^2^=0.879; Functional gene: P=0.001, R^2^=0.845; 16S-BC: P=0.001, R^2^=0.0.787; 16S-WUF: P=0.001, R^2^=0.726), and was the only significant variable for MASH (Figure 2) and 16S rRNA gene-based measures of beta-diversity (Figures S6, panels B and C) among those parameters evaluated. The functional gene ordination was also correlated, albeit weakly, to total 5-day precipitation (P=0.028, R^2^=0.359; Figure S6A). In order to control for spatial variance, a more rigorous db-RDA (30) was used on constrained NMDS ordinations, which allows the influence of a matrix of conditioning variables (i.e., location) to be “removed” prior to analysis. No significant associations (P>0.05) were found in the functional gene and OTU Bray-Curtis ordinations, however, the MASH and OTU weighted UniFrac distances were significantly associated with sampling time (ANOVA: F=1.274, P=0.031; F=2.174, P=0.04, respectively).

**Figure 2:**
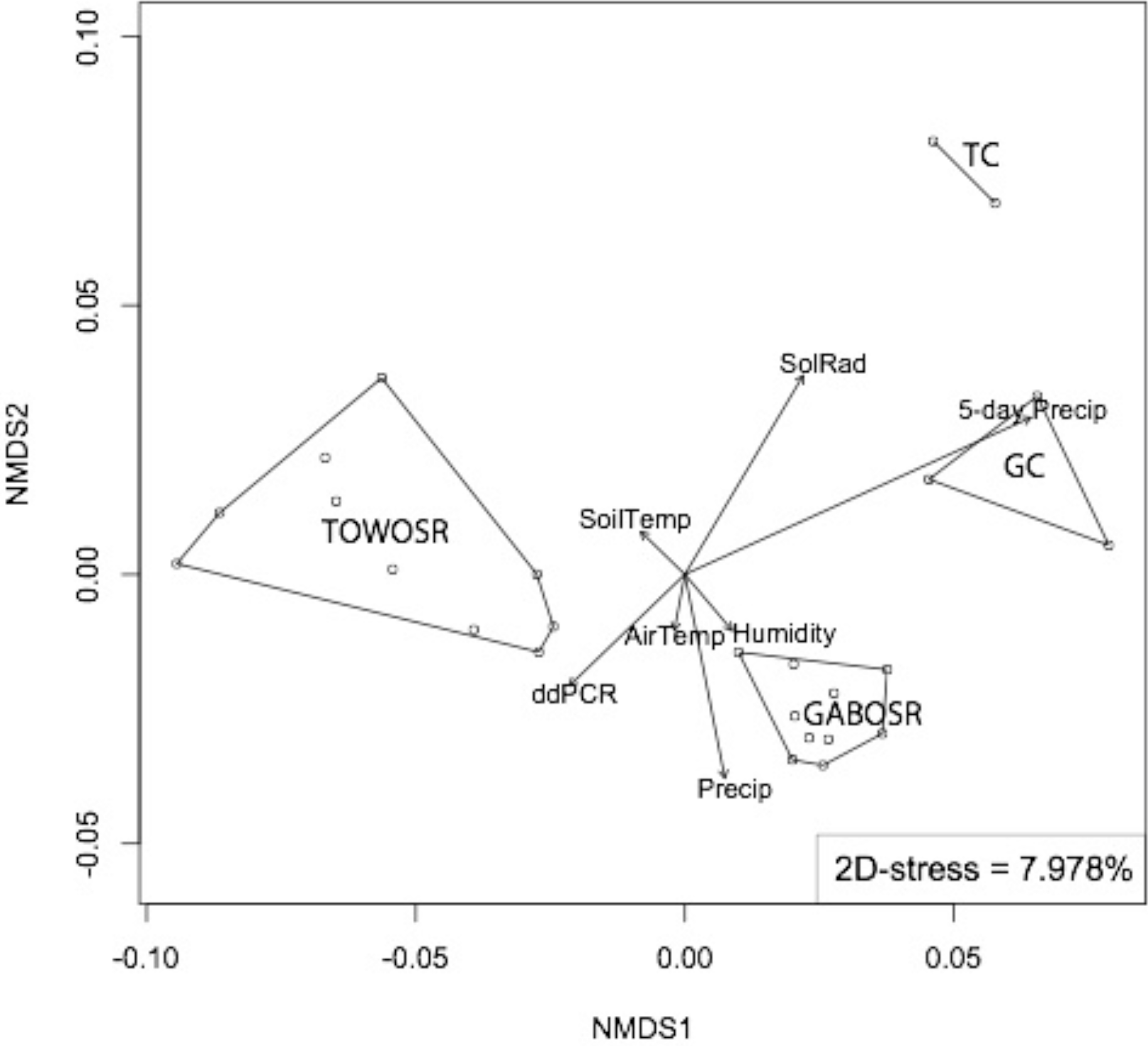
The effect of environmental parameters on microbial community structure. The graph shows non-metric multidimensional scaling (NMDS) of the sequenced communities based on whole-community MASH distances. Each dot represents a metagenome sample and those that were more similar to each other are grouped together by connected lines. Arrow vectors indicate correlation to metadata parameters.

### Culture-based detection of *E. coli* does not correlate with metagenome-based results

The abundance of *E. coli* in the metagenomes was low for all samples (~0.002% of total reads). Samples with the highest relative abundance of metagenomic reads matching to *E. coli* were negative for all culture-based tests (Table S3), which indicated spurious *in*-*silico* results (e.g., reads from non-*E. coli* genomes matching to conserved genes such as the rRNA operon). In addition, when using imGLAD (31) to predict the probability that *E. coli* was present in the metagenomes, a tool developed by our team to deal with spurious matches, all samples yielded a P-value of 1 (i.e., 0 probability of presence), which suggested that any *E. coli* populations (including STEC) were below the estimated limit of detection for the datasets in hand (3% coverage of *E. coli* genome at a minimum of 0.12 sequencing depth). The absolute abundance of the STEC based on ddPCR was also low (~1 in 10^8^ cells, assuming average molecular weight of a bp of DNA is 660g/mol, 5 Mb genome size, and 1 copy *stx*/genome) or absent in all samples, which supports our bioinformatic approaches (Table S3).

### Differentially abundant (DA) functions and taxa between locations

Of the 1,105 SEED subsystems (pathways) and 1806 taxonomic groups identified, 911 and 408 were significantly DA with P_adj_ < 0.05 for subsystems and taxa, respectively. Using pairwise comparisons between GABOSR, TOWOSR, and upstream sites, 184 SEED subsystems had Log_2_ fold change (L2FC) > 1, while 273 taxa had L2FC > 2, which were grouped into 36 and 35 broader functional and taxonomic categories, respectively (as described in the supplementary data). This analysis revealed several notable trends that were consistent between the SEED and taxa results (Figures 3 and S7). More specifically, iron acquisition genes appeared to more abundant in the upstream samples, particularly in the samples collected upstream of TOWOSR (TC1 and TC2). Plant-associated and photosynthesis genes were more abundant in the more pristine samples (WS1 and WS2). Consistently, members of the phyla, *Alphaproteobacteria* (e.g. *Rhizobiales*; see Supplementary data file S2), were more abundant upstream. The upstream sites were also DA for taxa that are associated with soil and aquatic habitats (e.g. *Gemmatimonadetes* and *Armatimonadetes*), which indicated that these sites may indeed receive less anthropogenic inputs, as we hypothesized.

**Figure 3:**
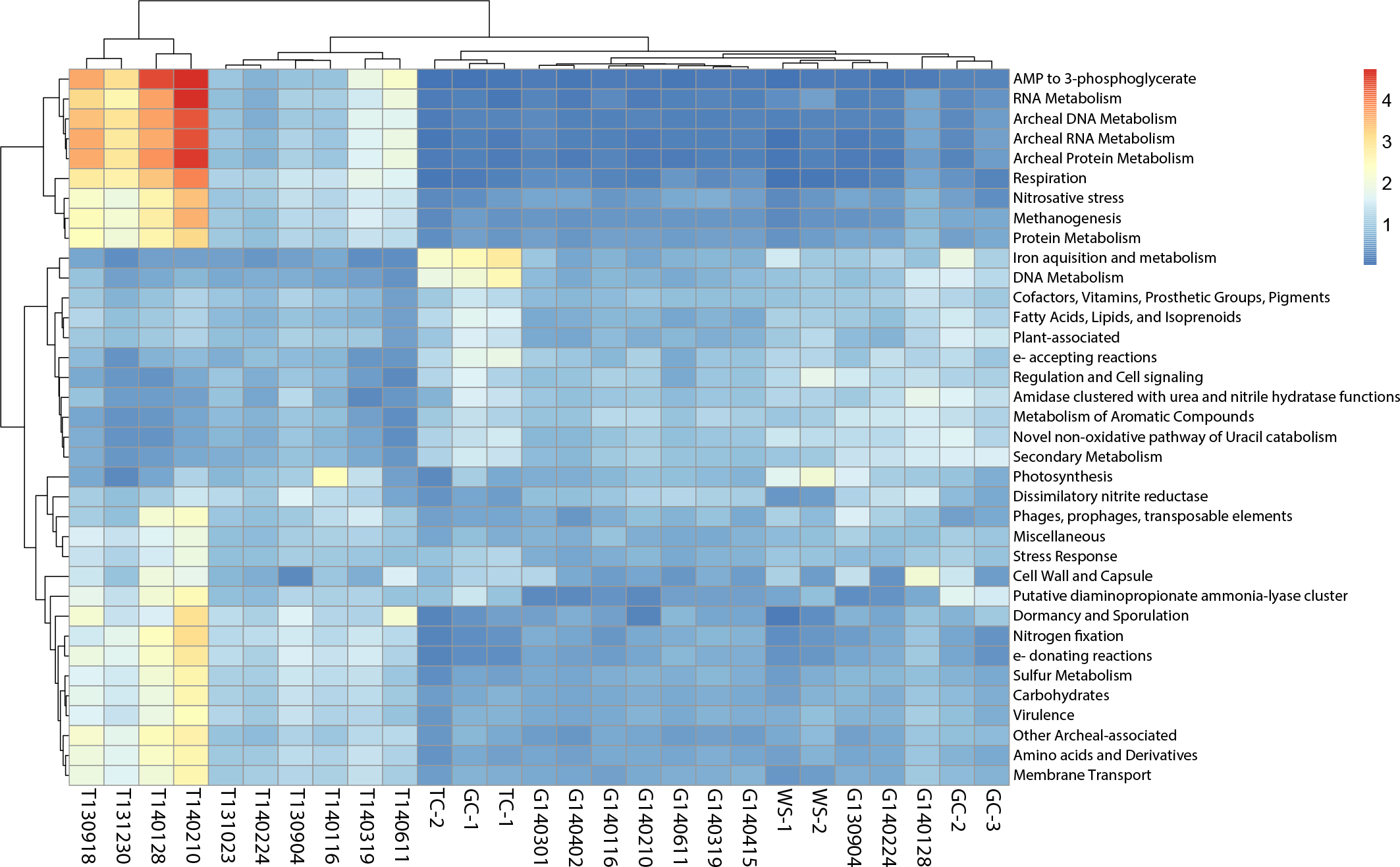
Functional profiles of creek sediment microbial communities. The heatmap shows SEED subsystems that were differentially abundant between locations (TOWOSR, GABOSR, and the upstream controls) with P_adj_ < 0.05. Color scale indicates the abundance relative to the average of all samples (increasing from blue to red).

Sample T140116 was enriched for both cyanobacteria based on OTU analysis (Figure S7) and photosynthesis genes (Figure 3). TOWOSR appeared to be DA in genes for anaerobic processes like anoxygenic photosynthesis and methanogenesis, along with genes related to archaeal DNA, RNA, and protein metabolism (all organisms known to carry out methanogenesis are *Archaea*). Consistently, the two TOWOSR samples (T140128 and T140210) that were most DA for archaeal and methanogenesis genes were also the most DA in *Archaea* and methanotrophs from the order *Methylococcales*, relative to the other sites. Other DA genes associated with anaerobic metabolisms, such as anoxygenic photosynthesis and sulfur metabolism genes (Figure 5), were congruent with taxonomic results that showed anoxygenic photosynthetic phyla *Chlorobi* (Green S bacteria), *Chloroflexi* (Green non-S), and the family *Chromatiaceae*, as well as known sulfur-metabolizing and anaerobic groups (e.g. *Thiobacillus* and *Clostridia*) to be more prevalent in the TOWOSR samples (Figure S7). Additionally, the TOWOSR samples, in general, were more abundant in the gut-associated phyla, *Firmicutes* and *Bacteroidetes*. Sample T140210 from TOWOSR was particularly enriched in specific enteric taxa: *Endomicrobia* and *Fibrobacteres*, which are rumen bacteria associated with cellulous degradation.

Collectively, these results suggested that our annotation and grouping methods were robust and that TOWOSR samples are more anaerobic, which could potentially indicate greater runoff and eutrophication as a result of human activity at this location. Also, the upstream sites were all significantly DA in *Actinobacteria* (i.e., common soil microbes and antibiotic producers), which provides further evidence in support of this system being a natural (and substantial) source of ARGs (see below).

### Quantifying anthropogenic and agricultural inputs

#### ARGs are more abundant in California samples compared to other similar environments

The abundance of ARGs in each dataset was determined by blastp search against the Comprehensive Antibiotic Resistance Gene Database (CARD; (32)). The most abundant ARGs detected are shown in Figure S8. A comparison of selected metagenomic datasets that included: metagenomes from agricultural sediments from Montana (MT) and soils from Illinois (Urb, Hav), more pristine/remote samples from the Kalamas River (Kal) and Alaskan permafrost (AK), as well as a highly polluted sample from the Ganges River (Agra), was performed in order to benchmark the level of anthropogenic signal observed in the Salinas Valley against other environments. The abundance of ARGs in the California samples were significantly greater compared to the other environmental metagenomes included here (Kruskal-Wallis χ^2^ =19.44, P = 0.0002; Figure 4A).

**Figure 4:**
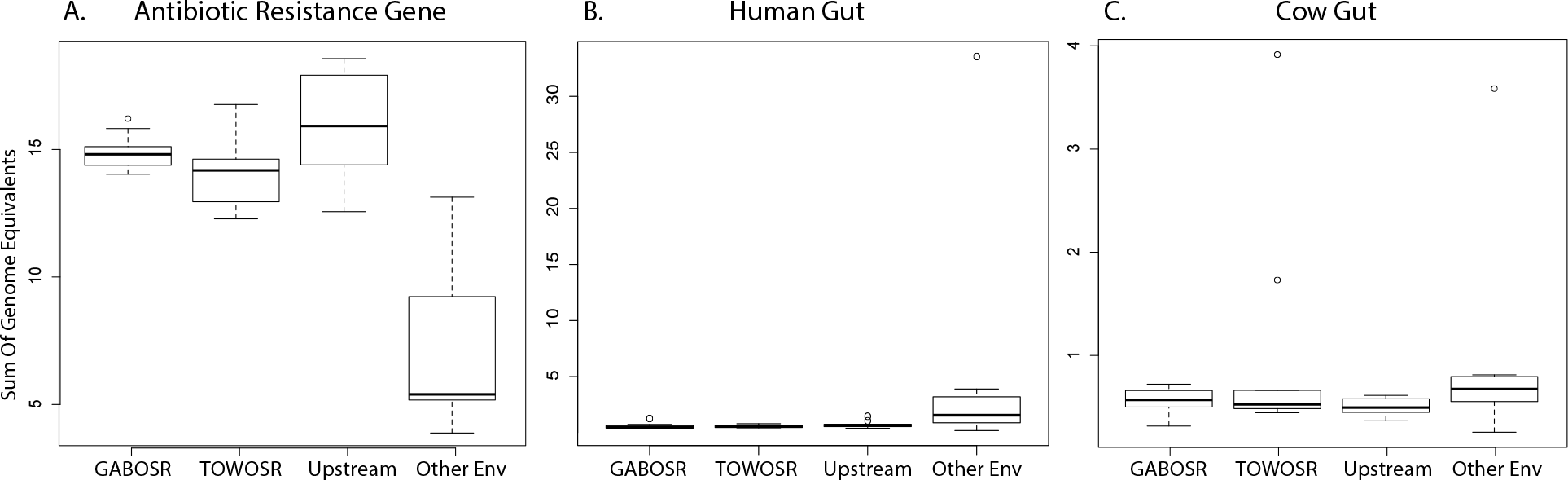
Abundance of ARG, human gut (HG), and cow gut sequences in the Salinas Valley metagenomes compared to other environmental metagenomes. The box and whisker plots show the interquartile range for the abundances with open dots indicating samples that exceeded 1.5× the interquartile range. The other environmental metagenomes (Other Env) included: 3 river sediments, 2 agricultural soils, 1 permafrost soil, and 2 river water samples from the Kalamas and Ganges Rivers.

**Figure 5:**
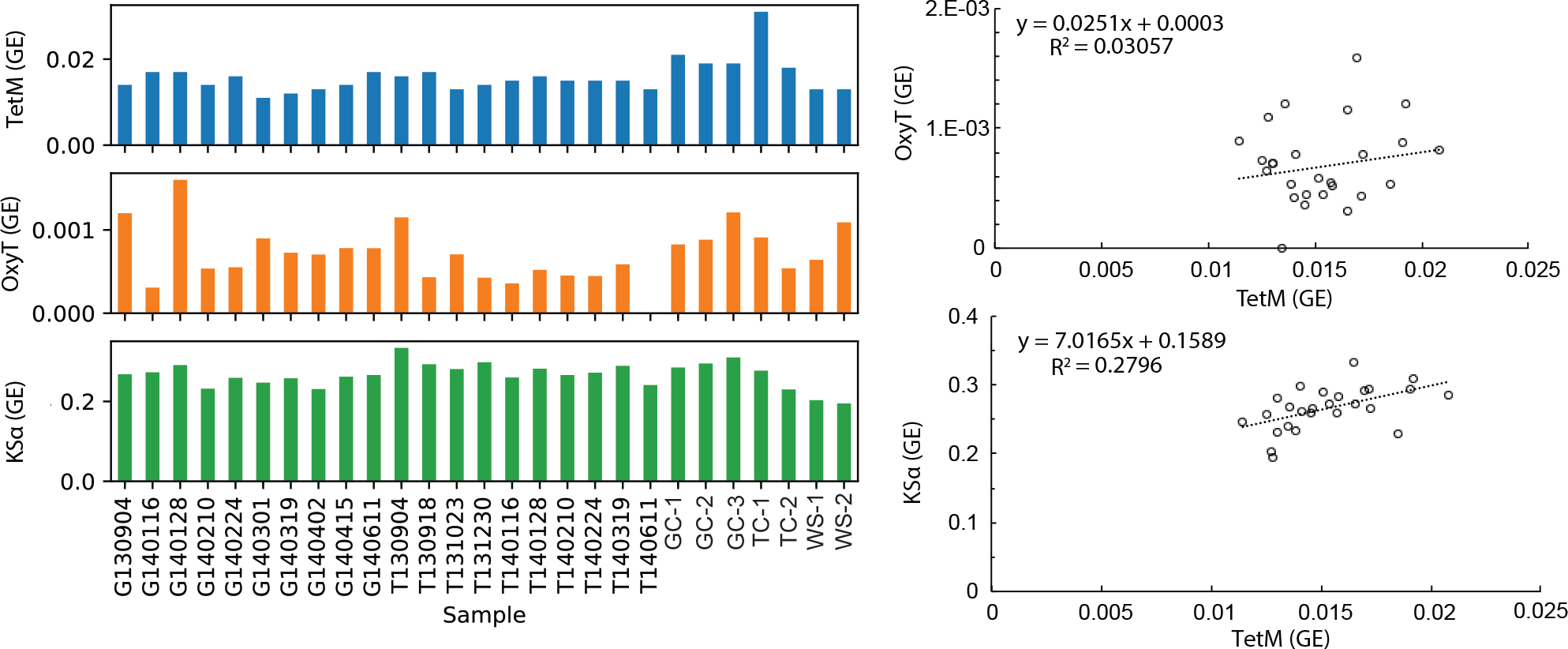
Abundances of selected antibiotic resistance and production genes in the Salinas Valley metagenomes. (**LEFT**) Abundance (expressed as genome equivalents) of *tetM*, *oxyT*, and *KSα* genes for the 27 sites included in this study. (**RIGHT**) Linear regression of *tetM* versus *oxyT* or *KSα* gene abundances. TC1 was an outlier for *tetM* abundance and was removed from this analysis.

#### Abundance of genes associated with antibiotics used in cattle

In order to better assess the impact (if any) of ARGs related to cattle ranching, we built ROCker models, a more accurate approach for finding metagenomic reads encoding a target gene of interest compared to simple homology searches (33), targeting tetracycline resistance (*tetM*) and production gene (*oxyT*) since tetracyclines are among the most common antibiotics used in livestock (34). We also built a model targeting ketosynthase alpha subunit genes (*KSα*), which are involved in the synthesis of many antibiotics, including tetracyclines (35). In order to exclude the effect of potentially confounding variables, only the California samples were used for linear regression analysis of the abundances of antibiotic production and resistance genes. ROCker analysis showed high prevalence of *tetM* in all samples and an abnormally high abundance for *tetM* was observed in sample TC1 (Figure 5, left panel). TC1 was thus considered an outlier and excluded from the linear regression analysis. The high abundance in TC1 was attributed to the fact that *tetM* has the widest host range of all tetracycline resistance (*tet*) genes due to its association with highly mobile conjugative transposons that behave similarly to plasmids and have several antirestriction systems (36, 37). *OxyT* did not significantly correlate to *tetM* abundance (r^2^ =0.031); however, *KSα* showed a moderate correlation to *tetM* (r^2^=0.280) (Figure 5, right panel).

#### Abundance of cow and human gut (HG) microbiomes

The signal from the Ganges River (Agra) sample greatly exceeded all other samples in both the absolute number (Table S4) and relative abundance expressed as genome equivalents (GE), i.e., the fraction of total genomes encoding human gut genes assuming a single-copy of each gene per genome (33.5 GE; 8-100× more abundant than all other samples; Figure 4B). There was a significant difference between the HG abundance averages observed in California metagenomes and the 8 metagenomes from 5 other habitats evaluated here (Kruskal-Wallis P=0.015). However, after correcting for multiple comparisons, none of the groups were significantly different (Wilcoxon Rank Sum P >0.1). Within California samples, there was no significant difference, overall, between abundances observed in the downstream samples and the average abundances of the upstream control samples (Kruskal-Wallis P=0.169).

The abundance of different cow gut genes had a similar trend to the human gut data (Table S4). However, two samples from TOWOSR (T140210 and T140611) showed an elevated signal for cow sequences (Figure 4C). Despite these two samples from TOWOSR with a higher level of cow gut signal, the average gene abundances were similar for California samples overall, and no significant difference was detected between the means compared to the other environmental metagenomes and the seven upstream control samples (Kruskal-Wallis P=0.090; Figure 4C).

## DISCUSSION

Analyses of river planktonic communities over time and land use have shown that these communities vary by average genome size, location, amount of sunlight, and nutrient concentrations (38) as well as by sampling time more so than space (39). However, the results presented here suggested that community composition of Salinas Valley creek sediments are structured primarily by spatial separation, and the local weather parameters tested here did not have a significant effect (Figure 2). More detailed *in*-*situ* metadata than those obtained here such as nutrient concentrations (e.g. organic carbon and biological oxygen demand) are needed in order to discern the processes that are driving community diversity and structure within each Salinas Valley site. For example, anaerobic taxa and processes related to methane and sulfur metabolism and anoxygenic photosynthesis were significantly more abundant in TOWOSR (Figure 3 and supplemental material), which suggests higher eutrophication from agricultural run-off or higher primary productivity by phototrophs, which was not reflected by the local weather parameters measured.

We compared abundances of reads annotated as ARG, human or cow gut in order to assess levels of anthropogenic impacts on Salinas Valley creek sediment communities. No significant difference was detected between the downstream samples and the upstream controls for any of the three anthropogenic indicators (Figure 4), which suggested that the land use practices surrounding the creeks does not have a lasting impact on the natural community and the inputs are likely diluted or attenuated faster than the intervals sampled here. We then benchmarked abundances observed in the creek sediments from this study against metagenomes from other environments. These included agricultural sediments and soils, permafrost, and river water from both pristine and polluted habitats. GABOSR, TOWOSR, and the upstream samples all had significantly higher ARG abundances compared to the average of the other environments tested here (Figure 4A). This high background level of reads annotated as ARGs suggested that the Salinas Valley creek sediments are a natural reservoir for these genes. Furthermore, resistance genes to synthetic antibiotics such as florfenicol (*fexA* and *floR*) and ciprofloxacin (*qnrS*), one of the most widely used antibiotics in humans worldwide, were absent or detected in very low abundance (less than 10 reads matching) in our datasets. Spurious matches to conserved gene regions can occur when analyzing short reads like the ones here, but the signal was not large enough to warrant further investigation using precise and targeted methods (e.g. ROCker). Overall, the absence of resistance genes to more recently introduced, synthetic antibiotics provides further evidence that the ARG signal observed in the Salinas Valley is likely autochthonous in origin. Future studies could involve deeper sequencing (higher community coverage) in order to recover long contigs and thus, determine the genomic background of the ARGs and if they are associated with mobile elements or plasmids for improved public health risk assessment. Still, these results highlight the importance of having a baseline or “pristine” sample to discern anthropogenic from naturally-occurring ARGs and have important implications for monitoring the spread of ARGs in the environment. For instance, without the upstream control samples, this study could have (speciously) concluded that GABOSR and TOWOSR are elevated in ARGs as a result of cattle ranching. However, the similar abundances found in the upstream samples indicated that the signal detected downstream could be inherent to this environment and that a more targeted analysis of specific ARGs was required to determine if the effect of cattle could be detected.

Tetracycline resistance genes have been shown to increase with and correlate to anthropogenic inputs along a river estuary system (40), suggesting that they can be useful indicators of anthropogenic pollution. However, tetracycline resistance genes are also found in other pristine or natural environments (28, 41–43), and therefore can also be considered part of the autochthonous gene pool in some habitats. Here, we tested the hypothesis that if tetracycline resistance genes are naturally occurring, the production enzymes for tetracycline should also follow similar abundance patterns, as antibiotic resistance and biosynthesis genes are often encoded on the same operon to ensure antibiotic-producing species are resistant to the product they synthesize (44). Thus, we expected to see a correlation between abundances of the tetracycline resistance gene, *tetM*, and its associated production genes (*oxyT*, *KSα*) if this system is not under heavy selection pressure of human-introduced antibiotics. The abundance of *tetM* in the Salinas Valley creek sediments was not correlated to *oxyT* and only moderately correlated to *KSα* (Figure 5). *OxyT* had very low abundance (less than 8 reads matching per sample), which suggested that the lack of correlation to *tetM* could be due to database limitations. That is, only a few reference *oxyT* genes are publicly available (13 sequences) and these likely do not capture the total diversity of this gene found in the environment. *KSα*, on the other hand, represents a broad class of synthesis genes for many different antibiotics with many more sequences in the reference databases and thus, a better estimate of antibiotic production potential was obtained based on these genes. Overall, these findings further supported that this ecosystem is a natural reservoir for ARGs, and the presence of tetracycline resistance is not likely to be caused by inputs from the cattle ranches. However, future investigations could involve additional antibiotic production gene references for more robust conclusions.

When compared to the other pristine or rural environmental metagenomes such as agricultural sediments and soils, permafrost, and river water, the abundances of reads annotated as human gut in the California sediments were not significantly different overall. However, the Ganges River (Agra) sample, collected from one of the most densely populated and highly polluted areas surrounding the river (Agra, Uttar Pradesh, India), was 1-2 orders of magnitude more abundant for human gut, compared to the rest of the samples used in our study (Figure 4B). Thus, a high human gut signal was expected for the Ganges River, consistent with previous results (45) and served as a reference to assess relative levels of human fecal contamination. The rest of the samples included in our comparisons were from rural/agricultural or more remote areas, with lower population density, and consistently had lower signals of human fecal contamination than the Agra sample. Therefore, the abundances of human gut sequences observed in Salinas Valley were consistent with the lower levels of human activity/density input relative to the other sites used for comparison here and indicated that our annotation and filtering methods were robust. Collectively, these results showed that metagenomics of river/creek sediments provide a reliable means for assessing the magnitude of the human presence/activity, consistent with recent studies of other riverine ecosystems (39, 45).

Contrary to the results for human gut, the abundances of cow gut signal in the California samples were not consistent with our expectations. The TOWOSR and GABOSR sites are directly downstream of large cattle ranch operations and identical pathogen recovery from water and upstream cattle indicated the cattle ranches were the source of fecal contamination (1). As such, we expected to see a higher level of cow signal in the downstream metagenome samples, yet the abundance was not significantly different from the other environments or the upstream controls (Figure 4B&C). Notably, two of the samples from TOWOSR (T140210 and T140611) showed elevated signal for cow that was similar to the abundance observed in the highly polluted Ganges River reference metagenome (Figure 4C). These samples (especially T140210) had a higher abundance of the rumen enteric and cellulose degrading taxa (*Endomicrobia* and *Fibrobacteres*; Figure S7), which supports the conclusion that these samples contained run-off from cattle, however the signal might be patchy or muted in the sediment and require more frequent sampling and/or larger sampling volumes than those used here to detect these signals.

Additionally, we were unable to detect any *E. coli* populations in any of the metagenomes, including samples that were positive for STEC via enrichment culture, indicating that it is not an abundant member of the sediment community (Table S3). This was consistent with imGLAD estimates that the sequencing effort applied to our metagenomes imposed a limit of detection for *E. coli*, and ddPCR results that showed abundance of STEC was low or absent in all samples. Overall, these results suggested that using shotgun metagenomics may not be sensitive (or economical) enough as a monitoring tool to detect a relatively low abundance microorganism in lotic sediments at the level of sequencing effort applied here, which was insufficient partly because of the extremely high community diversity (Figure S1). More than the 2.5 to 5 Gbp/sample sequencing effort applied in this study would have been required to detect ~10 *E. coli* cells in a sample according to our estimates, which is not economical based on current standards and costs. More specifically, obtaining the imGLAD minimum threshold of 0.12× coverage for an STEC genome (5 Mbp) in our metagenome libraries (average 4 Gbp), would require 0.6 Mbp of STEC reads, or 0.015% of the total metagenome, which translates to a relatively large number of cells *in situ*. For example, assuming 10^8^ total cells/g of sediment, it would require ~10^4^ STEC cells/g of sediment to robustly detect in the metagenomes (or 100 times more sequencing for detecting ~10 cells/g). Thus, the limit of detection of metagenomics, as applied here, was not low enough and should be combined with methods that offer lower detection limits and more precise counts (such as ddPCR).

Rivers are highly dynamic ecosystems and therefore subject to higher random variation and sampling artifacts that likely affect the dilution of the exogenous (human) input. Further, our samples represent relatively small volumes of sediment (~10 g) and the resulting metagenomic datasets did not saturate the sequence diversity in the DNA extracted from these samples (Figure S1), which might introduce further experimental noise and stochasticity. Despite these technical limitations, our data consistently showed little evidence that agricultural or cattle ranching activities have a significant effect on the creek sediment microbial communities. The underlying reason for these results remains speculative but should be the subject of future research in order to better understand the impact of these activities on the environment. Additionally, the level of functional and taxonomic diversity, as well as the sample heterogeneity (especially in TOWOSR), suggested that shorter intervals between sampling as well as *in situ* geochemical data are needed to elucidate the fine scale processes driving the community composition within each location. Although the continued presence of STEC in Salinas watershed sediments is a public health risk, we did not find evidence that runoff from human activities has a substantial effect on the sediment microbial community when compared to more pristine sites. An imperative objective for public health is to assess how and where current agricultural practices impact the environment in order to determine best practices. The work presented here should serve as guide for sampling volumes, amount of sequencing to apply, and what bioinformatics analyses to perform on the resulting data for future public health risk studies of river water and sediment habitats. Finally, the ROCker models developed here for tetracycline resistance and production genes should be useful for robustly examining the prevalence of these genes in other samples and habitats.

## Acknowledgements

This work was supported by the USDA (award 2030-42000-050-10), the US National Science Foundation (awards No 1511825 and 1831582 to KTK) and the US National Science Foundation Graduate Research Fellowship under Grant No. DGE-1650044. The funding agencies had no role in the study design, data collection and analysis, decision to publish, or preparation of the manuscript.

## MATERIALS AND METHODS

### Sample collection and enrichment method for STEC

Sediment samples were collected from watersheds at public-access locations (Table S1). Weather information was downloaded from the California Irrigation Management Information System database (http://ipm.ucanr.edu/calludt.cgi) for the day of and five days prior to the sampling day from the closest monitoring station to the downstream sites (Table 1). Sediment was suspended into the water column using a telescoping pole and approximately 250 mL of sample (suspended sediment and water) was collected immediately in a sterile bottle. All samples were transported on ice and processed within 24 hours. DNA from 10 g of resuspended sediment/water mix was purified for sediment DNA using MoBio PowerSoil DNA extraction kit, following the manufacturer’s protocol. A separate 100 mL of the sample was used for enrichment and isolation of STEC as previously described (15).

### PCR-based quantification method for STEC

Droplet digital PCR (ddPCR, BioRad) was performed on sediment DNA following the method of Cooley et al. (19). Each 20 μL reaction used 10 μL BioRad’s Supermix for Probes, 2 μL primer (0.3μM final concentration) and probe (0.2μM), up to 1 μg DNA, 1.2 μL MgCl2 (1.5mM), and 0.2 μL HindIII (0.2 U/μL). Primer and probe sequences were as previously published for STEC (19). Droplets were created with Droplet Generation Oil for Probes in the QX-200 droplet generator (BioRad), and amplified for 5min at 95°C, 45 cycles at 95°C for 30 s and 60°C for 90 s, then 5min at 72°C and 5min at 98°C. Droplets were processed with the QX-200 Droplet reader and template levels were predicted by QuantaSoft software version 1.7.4 (BioRad).

### DNA sequencing and Bioinformatics sequence analysis

#### Metagenomic sequencing and community coverage estimates

Shotgun metagenomic sequencing libraries were prepared using the Illumina Nextera XT library prep kit and HiSEQ 2500 instrument as described previously (46). Short reads were passed through quality filtering and trimming as described previously (47). Average community coverage and diversity were estimated using Nonpareil 3.0 (29) with kmer kernel and default parameters. Sequences were assembled with IDBA (48) using kmer values ranging from 20 to 80.

#### Taxonomic analysis of rRNA gene-encoding sequences

Metagenomic reads encoding short subunit (SSU) rRNA genes were extracted with Parallel-Meta v.2.4.1 using default parameters (49). Closed reference OTU picking at 97% nucleotide identity with taxonomic assignment against the GreenGenes database (19) was performed using MacQiime v.1.9.1 (51) with the reverse strand matching parameter enabled and the uclust clustering algorithm (52). Alpha diversity was calculated as the true diversity of order one (equivalent to the exponential of the Shannon index) and corrected for unobserved species using the Chao-Shen correction (53) as implemented in the R package entropy (54). Richness was estimated using the Chao1 index (55), and evenness was calculated from the estimated values of diversity divided by richness. Significant differences in taxonomic diversity, evenness, and richness were assessed using two-sided t-tests. Multiple rarefactions were performed on OTU tables as implemented in MacQiime v.1.9.1 (rarifying up to the minimum number of counts per sample: option -e 5,596).

#### Determination of the total community bacterial fraction

In order to determine whether bacterial gene abundances need be corrected for relative bacterial fraction in the total metagenome libraries, the relative abundance of *Bacteria*, *Archaea*, and *Eukarya* was estimated in each dataset by searching a subset (~1×10^5^ reads per sample) of randomly selected protein coding reads against the TrEMBL database ((56); downloaded May 2018) using DIAMOND blastx v.0.9.22.123 (57) with the “--more sensitive” option and e-value cutoff of 1×10^−5^. The TrEMBL IDs for best hit matches were summarized at the domain level using custom scripts and the metadata files available at ftp://ftp.uniprot.org/pub/databases/uniprot/current_release/knowledgebase/taxonomic_divisions/ No significant difference in the relative abundance of *Bacteria* was found between the different samples, thus no correction for bacterial fraction was applied to gene abundance calculations.

#### Functional and ARG annotation of metagenomic sequences

Protein prediction was performed using FragGeneScan adopting the Illumina 0.5% error model (58). Resulting amino acid sequences were searched against the Swiss-Prot (downloaded June 2017) (56) and Comprehensive Antibiotic Resistance gene (CARD, downloaded May 2017; 26) databases using blastp (59) for functional annotation. Best matches to the Swiss-Prot database with >80% query coverage, >40% identity and >35 amino acid alignment length were kept for further analyses. A more stringent cut off was used for best matches to the CARD (>40% identity over >90% of the read length) to minimize false positive matches.

#### Detection of cow and human gut microbiome associated sequences

Searches for cow gut associated sequences were performed using our own collection of cow fecal metagenomes from six cow individuals collected in Georgia, USA. DNA extracted from cow fecal material underwent the same library prep, DNA sequencing and quality trimming and processing as described above. Short reads for both the cow gut and CA sediment metagenomes have been deposited to the SRA database (submission IDs: PRJNA545149 and PRJNA545542, respectively). Predicted genes (as nucleotides) from all six individual cows were pooled together and de-replicated at 95% identity using the CD-HIT algorithm (Options: -n 10, -d 0; (60)) resulting in 459,176 non-redundant cow gut metagenome “database” sequences. Human gut-associated sequences were assessed based on comparisons of short-reads against the Integrated Gene Catalog (IGC) of human gut microbiome genes (61), heretofore referred to as Human Gut Database (HG) for clarity. The abundance of cow and human gut signal in the short-read metagenomes was determined based on the number of reads from each dataset matching these reference sequences using blastn v2.2.29 with a filtering cut off of >95% identity and >90% query length coverage.

#### Abundance of specific antibiotic resistance (ARG) and production genes using ROCker

Dynamic filtering cut-off models targeting a tetracycline resistance gene (*tetM*) and two antibiotic production genes (*oxyT* and *KSα)* were designed with ROCker v1.3.1, as previously described (33). Reference sequences for model building were manually selected from public databases and models were built for 150bp reads and default parameters. The reference sequences and ROCker models are available at http://enve-omics.ce.gatech.edu/rocker/models. Short reads were searched against the reference sequences used to build the model with blastx. The ROCker models were used to filter matches, which were subsequently divided by the median reference gene length in order to calculate sequencing coverage and were then normalized for genome equivalents as described below. Correlation between abundances of antibiotic production and resistance genes was determined using linear regression.

#### Quantification of genome equivalents (GE)

Average genome size and genome sequencing depth (i.e., the average sequencing depth of single copy genes) were determined for each sample using MicrobeCensus v1.0.6 with default parameters (62). The sequencing depth of reference genes with a given annotation was estimated for each dataset (in reads/bp), then divided by the corresponding average genome sequencing depth and summed to give the total GEs per sample.

#### Mash and multivariate analysis

MASH v1.0.2 (63) was used to assess overall whole-community similarity among metagenomes in a reference database-independent approach (Options: -s 100000). Functional gene and 16S rRNA gene-based OTU count matrices were median-normalized using the R package DESeq2 (v.1.16.1; (64)). Pairwise Bray-Curtis and weighted UniFrac (16S only) dissimilarity indexes of the normalized counts were used for principal component analysis (PCA) and non-metric multidimensional scaling (NMDS) analysis in order to assess whole-community gene functional and taxonomic (16S rRNA gene OTUs) similarity. The significance of metadata parameters on the NMDS ordinations was performed using the ecodist and envfit functions of the R package vegan v2.4.4 (indices included: location, sampling time, ddPCR counts for STEC, same day precipitation, 5-day precipitation, solar radiation, air temp, soil temp, and humidity). The two west Salinas samples (WS1 and WS2) were excluded from this analysis in order to minimize confounding variation of temporal and spatial differences. In order to control for spatial variance, a more rigorous distance-based redundancy analysis (db-RDA; (30)) was used to investigate the correlation to metadata using the capscale function in the R package vegan (included same indices as above, but with Condition(location) constraint on ordinations).

#### In-silico detection of E.coli in sample metagenomes

The presence of any *E. coli* in the metagenomes was determined using a blastn search of short reads against an STEC reference genome (accession NC_002695) that had been filtered to remove non-diagnostic (i.e. highly conserved among phyla) regions with MyTaxa (65). Only matches with nucleotide identity >95% and alignment length >97% were used to calculate relative abundance of *E. coli* in the metagenomes. This level of sequence diversity (nucleotide identity >95%) encompasses well the diversity within the *E. coli*-*Shigella spp*. group; thus, any *E. coli* populations present in the metagenomes at high enough abundance would be detected at this filtering cutoff. The best hit output from blastn was also analyzed with imGLAD (31), a tool that can estimate the probability of presence and limit of detection of a reference/target genome in a metagenome.

#### Determination of differentially abundant (DA) taxa and gene functions

Functional annotations of the recovered protein sequences were summarized into several hierarchical ranks including metabolic pathways and individual protein families based on the SEED classification system (66). The 16S rRNA gene OTUs were placed into taxonomic groups based on the lowest rank of taxonomic classification (genus, family etc.) shared by 90% or more of the sequences within the OTU using MacQiime v.1.9.1 (51). DA functional annotation terms (subsystems) or OTUs were identified in samples grouped by location (e.g., pairwise comparison of all 10 TOWOSR vs. all 10 GABOSR and vs. all 5 upstream “pristine control” sites) using the negative binomial test and false discovery rate (P_adj_ <0.05) as implemented in DESeq2 v1.16.1 (64). Subsystems with Log_2_ fold change (L2FC) >1 or taxa with L2FC >2 were manually grouped into broader categories based on known functional or taxonomic similarities, respectively (Figures 3 & S7), which were then normalized by library size (per million read library). A larger L2FC cutoff was used for taxa to account for the larger dataset size and allow for inspection of the taxa contributing most to differential abundance between the locations. The taxonomic assignment of these DA taxa were confirmed against the SILVA database (downloaded October 2018; (67)). Each subsystem or taxonomic category was then divided by its average sequencing depth across all samples to provide unbiased counts for presentation purposes.

#### Comparison of putative anthropogenic signals observed in California sediments to metagenomes from other environments

Publicly available metagenomes from other studies were used to compare abundances of reads annotated as ARG, HG, and cow gut with the results obtained for the California sediment datasets reported here. These metagenomes included: three Montana River sediments (MT; (20)), two temperate agricultural soils from Illinois (Hav and Urb; (68)), an Alaskan tundra soil (AK; (69)), one sample from the Ganges River near Agra, Uttar Pradesh (Agra; (45)), and one from the Kalamas River in Greece (Kal; (39)). Short read metagenomes for MT samples were downloaded from MG-RAST ((70); MG-RAST IDs: 4481974.3, 4481983.3, 4481956.3). The remaining datasets were obtained from the NCBI short read archive (SRA) database (Hav: ERR1939174, Urb: ERR1939274, AK: ERR1035437, Agra: SRR6337690, Kal: SRR3098772). Reads from these metagenomes were comparable to the ones from this study (100 – 150bp paired-end Illumina sequencing) and underwent the same trimming, annotation (against the CARD, HG, and cow gut databases only) and gene count normalization protocol as described above. The Kruskal-Wallis test in R was performed to determine significantly different mean abundances between groups. Alpha diversity and taxonomic comparisons were performed (for MT datasets only) based on metagenomic reads encoding fragments of the 16S rRNA gene, which were identified as described above.

